# Differential Speed and Accuracy Trade-off in Working Memory Retrieval and Bilateral Precuneus between Older Men and Women

**DOI:** 10.1101/2024.10.05.616832

**Authors:** Darlingtina K. Esiaka, Stephanie Strothkamp, Aghayeeabianeh Banafsheh, Lucas Broster, David K Powell, Gregory Jicha, Yang Jiang

**Author notes:** Corresponding Authors: Darlingtina Esiaka, PhD; Email Address; Center for Health, Engagement, and Transformation, University of Kentucky College of Medicine. 760 Press Avenue, Suite 426, Lexington, KY. 40536; Yang Jiang, PhD; Email Address; Department of Behavioral Science, University of Kentucky College of Medicine. 113 Medical Behavioral Science Building, Lexington, KY. 40536.

## Abstract

**Background:** Despite various hypotheses, including differences in longevity, hormones, genetics, and neuroanatomy, the reasons for the higher prevalence of Alzheimer’s disease in older women compared to men remain unclear. Emerging evidence suggests that the precuneus, a key region of the default mode network, is linked to internally focused processes like memory retrieval. This study examined sex differences in the relationship between precuneus volumes and working memory retrieval speed in cognitively normal older adults, hypothesizing that disparities in precuneus size and function contribute to reduced working memory performance in older women.

**Method:** A cohort of participants (N=45; 25 women; M_age_ =77) from the University of Kentucky Alzheimer’s Disease Research Center completed the Bluegrass Working Memory Task while undergoing 3T Siemens magnetic resonance imaging scans.

**Result:** Applying Spearman correlation analyses, the results revealed correlations between working memory accuracy and volumes in the left (r=-.43, p<.01) and right (r=-.36, p<.05) precuneus across all subjects. Sex difference analysis indicated a tendency for the accuracy of the memory task to correlate more frequently with the left precuneus in women (r= 0.54; p < 0.05) than in men. Similarly, volumes in the left precuneus displayed a significant negative correlation with reaction time in response to memory target (*r* = -0.426; p < 0.05) and memory distractor (*r* = -0.549; p < 0.01) in women. There is a sex difference in accuracy and speed trade-off. While men were faster in reaction time, women were better in the accuracy of the memory task. Particularly noteworthy was the consistent association in women, where neurocognitive measures (Trail A, r= -.50, p<.01; Trail B, r= -.06, p<.01) reliably correlated with volumes in the left precuneus—a relationship not observed in men.

**Discussion:** Our findings suggest that the left precuneus volume is associated with processing speed and accuracy of working memory performance, especially in women. Given that the left precuneus plays a key role in supporting various aspects of cognition, including memory retrieval, our findings point to the potential of reaction time serving as a surrogate marker for fMRI in predicting cognitive decline, particularly when considering sex differences.

## Introduction

Alzheimer’s disease (AD) is a progressive neurodegenerative disorder, associated with impairments in various cognitive functions, including working memory (Kirova et al. 2015). AD disproportionately affects women (Ferretti et al. 2018; Meilke et al. 2020), with approximately two-thirds of those affected by AD being women (Meilke et al. 2014). While the precise cause of sex differences in Alzheimer’s disease (AD) remains unclear, numerous studies have associated various factors with the observed distinctions between men and women in AD.

The examination of sex differences in AD often contemplates the longevity theory, given that women generally outlive men, making age a significant risk factor for AD (Alzheimer’s Association, 2024). Nevertheless, conflicting evidence exists regarding whether age-adjusted AD risk is higher in women. Some studies found no significant sex differences in AD incidence (Fiest et al., 2015; Neu et al., 2017), while others suggested higher rates in women (Altman et al., 2014), especially after the age of 80 (Ruitenberg et al., 2001; Roberts et al., 2014). Additionally, hormonal changes during menopause have been implicated in the elevated risk of AD in women. Research indicates that biomarker abnormalities associated with AD are most pronounced in postmenopausal women (Mosconi et al. 2017; Oveisgharan et al. 2018).

In addition, apolipoprotein ε4 allele (APOE4) has been identified as a factor contributing to AD risk between sexes, with women carrying APOE4 experiencing faster cognitive decline (Altmann et al., 2014; Mosconi et al., 2017). Moreover, sex differences in AD manifestation extend to women exhibiting more severe neuropsychiatric symptoms, neuroinflammation, and cognitive impairments (Cui et al., 2023; Eikelboom et al., 2022; Law et al., 2018). Significantly, structural brain differences and anatomical changes in mild cognitive impairment and AD have been observed, indicating distinct effects on men and women (Ardekani et al., 2016; Cavedo et al., 2018; Kim et al., 2015; Koran et al., 2017). These findings contribute to a deeper understanding of the complex nature of sex differences in AD, encompassing genetic, hormonal, neurocognitive, and pathophysiological factors.

A growing body of evidence points to the importance of regions outside the traditional focus of medial temporal lobe, hippocampus, particularly the precuneus, in predicting AD progression. The precuneus, situated in the posterior medial parietal cortex, plays a crucial role in visual-spatial image processing and episodic memory retrieval (Dadario & Sughrue, 2023; Dordevic et al., 2022; Eskildsen et al., 2015). Studies suggest that decreased precuneus volumes may serve as an indicator of AD pathology, independent of medial temporal lobe involvement (Billette et al., 2022; Casula et al., 2023; Haussmann et al., 2017; Mahady et al., 2021; Richter et al., 2022). Billette et al. (2022) used a novelty-related fMRI activity in medial temporal lobe regions and the precuneus to show that increased precuneus activity might be an early indicator of memory impairment. A 2017 study found that patients with aMCI demonstrated cortical thinning in the medial temporal cortex and precuneus (Haussmann et al., 2017).

It has long been reported that men have larger gray matter volumes in the left parietal lobe, particularly inferior parietal lobe. In contrast, in women are larger in the right inferior parietal relatively to the left parietal (Frederikse et al., 1999; 2000). The reason is not clear and under debate. Sex-specific differences in precuneus connectivity further contribute to the complexity of AD research (Williamson et al., 2022; Zhang & Chiang-shan, 2012), highlighting the need to consider both structural and functional aspects. Zhang and Chiang-shan (2012) observed sex differences in precuneus connectivity, with men exhibiting greater connectivity with the dorsal precuneus in the cuneus and medial thalamus, while women showed increased connectivity with the ventral precuneus in the hippocampus/parahippocampus, middle/anterior cingulate gyrus, and middle occipital gyrus compared to men. Similarly, Williamson et al. (2022) examined sex differences in the hippocampal connectivity to different areas of the brain and found that hippocampal connection to the precuneus cortex was significantly stronger in men than in women. Overall, despite various hypotheses, including differences in longevity, hormonal changes, genetics, neuroanatomy, and pathophysiology, the precise reasons for the higher prevalence of AD in older women compared to men remain unknown. Understanding these differences may offer valuable insights into developing targeted interventions for cognitive health.

In recent years, working memory—the cognitive system responsible for temporarily holding and manipulating information without emotional content, has emerged as a crucial domain in understanding AD risks (Borhani et al. 2021; Zheng et al. 2023). There is growing evidence that deficits in working memory may serve as an early indicator or a contributing factor to the development of cognitive impairments (Kjærstad et al., 2023). The intricate interplay between the integrity of working memory processes and the pathological changes observed in AD underscores the significance of investigating this link for both early detection and a deeper comprehension of the cognitive underpinnings of AD. However, there is a paucity of studies examining sex differences in non-working memory, particularly the significance of functional magnetic resonance imaging (fMRI) findings in conjunction with behavioral measures in healthy older adults.

Sex differences in working memory performance have been a subject of considerable research interest, with findings suggesting nuanced patterns. Some studies indicate that women exhibit advantages in verbal working memory tasks (Goldstein et al., 2005; Hirnstein et al., 2023; Voyer et al., 2021). Others find that men show advantages in spatial working memory tasks (Chen et al., 2020). Mixed findings on sex differences have also been reported regarding the accuracy, reaction times, and false alarm rates during memory performance. Subtle variations in these parameters have been observed between men and women (Chang & Moscovitch, 2022; Samson et al., 2023), with women outperforming men in immediate and delayed recall (Sundermann et al., 2016). However, Vaughan and Birney (2023) emphasize the importance of considering individual variability and the specific nature of the memory tasks employed. There remains a need for a more nuanced and multifaceted understanding of sex differences in working memory.

Determining if and where sex differences exist in AD is critical for the direction of future targeted diagnostics and therapeutic interventions. The current study aims to explore potential sex differences in memory performance and precuneus brain volume among cognitively normal older adults. Specifically, we examined the volume of the precuneus region involved in memory and compared participants’ performance on a memory task. Our hypothesized pattern is that sex differences would result in differential effects on precuneus volumes and network-associated working memory performance in cognitively normal older adults. By delving into the relationship between non-working memory and the risk of cognitive decline, we aim to shed light on how deficits or alterations in non-working memory may serve as potential predictors or indicators of cognitive decline, providing valuable insights for early detection of cognitive decline.

## Materials and methods

### Participants

We recruited fifty-two (52) cognitively normal older adults (*m*_age_ = 76.7 years; 27 women) from the University of Kentucky Alzheimer’s Disease Center cohort for the study. All of the participants were racialized as non-Hispanic White Americans. Seven cases were removed from the analysis based on the exclusion criteria—the presence of dementia, double participation in the study, and missing data on neuropsychological tasks. One participant had dementia, while six participants completed the test two times. Their second records, which also had more missing data in their behavioral task, were removed. A total of 45 participants were included (*M*_age_ = 77.05, *SD_age_* = 7.48; 25 women) in the present study. All research activities were approved by the University of Kentucky Institutional Review Board, and all participants provided written informed consent.

### Measures

#### Bluegrass Short-Term Memory Task

The Bluegrass Short-Term Memory Task (BeST) is an adapted version of a previous fMRI delayed match-to-sample memory task (Jiang et al., 2000) and previously described (Jiang et al., 2016). The task uses a 10-minute run time in which two pictures/images are encoded in each trial. Pictures of common images (e.g., candle, hammer, etc.) are used as the targets and distractors (i.e., match and nonmatch). In the BeST 10-min older-adult friendly version, two sample pictures were encoded for each given trial. This reduces scanning time for older adults and increases the number of matches with a balanced number of non-matches. Images are scrambled between trials to ensure the baseline of activity is reached. A representation of the task is shown in Figure 1.

**Figure 1.**
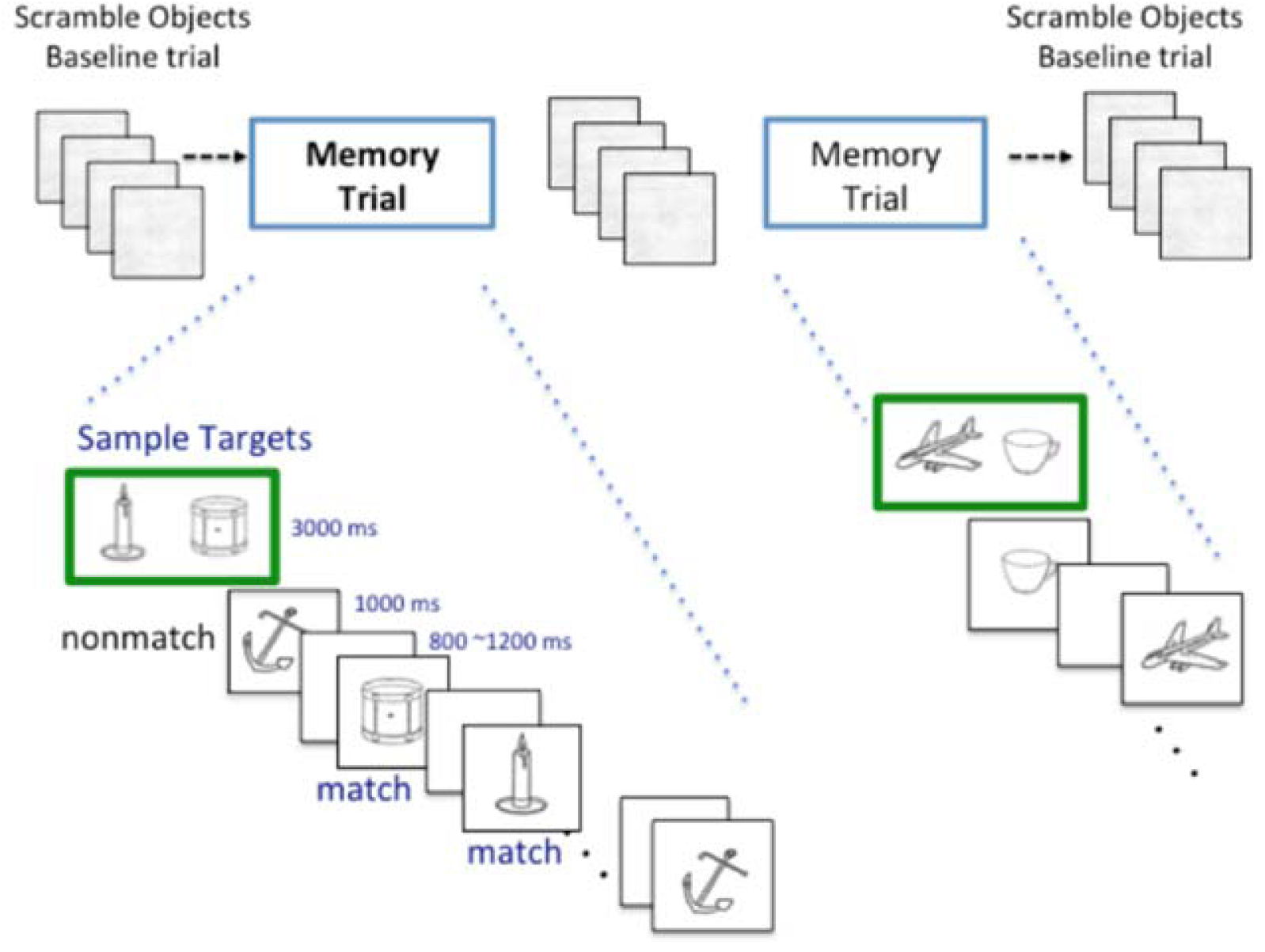
The Bluegrass Short-term Memory task (BeST) is described in Jiang (2016). In this task, participants memorize two sample images and then indicate whether each image presented thereafter matches either sample target image.

Memory performance was characterized by accuracy and reaction time (RT). We measured accuracy by the percent/ratio of correct answers and false alarms. A false alarm is the action of identifying a non-match visual object as a match target held in memory in a delayed match-to-sample paradigm. Participants are shown two target items to remember at the beginning of a trial and asked to determine whether or not other items seen in the trial are a match or non-match (e.g., Thinking a car in a parking lot is yours when it is not). Participants switched hands, indicating memory targets and non-targets (or distractors) between blocks. Thus, the RTs of memory retrieval of targets and distractors are balanced within a subject. In this study, we performed two trials of the BeST Memory Task for each participant, looking at the performance for both targets and distractors. False Alarm Rate (FA_Rate), which increases with cognitive aging, was also measured. Accuracy and RTs were measured for the first and second-trial match and non-match targets. False alarm number (FA_N), FA_Rate, and false alarm reaction time (FA_RT) were also measured.

#### Trail Making test

The Trail Making Test (TMT; Bowie & Harvey 2006) is a neuropsychological test that provides information on visual search, scanning, speed of processing, mental flexibility, and executive functions. The TMT consists of two parts. TMT-A requires an individual to draw lines sequentially connecting 25 encircled numbers distributed on a sheet of paper. Task requirements are similar for TMT-B, except the person must alternate between numbers and letters (e.g., 1, A, 2, B, 3, C, etc.). Participants were instructed to complete each part of the TMT as quickly and accurately as possible. The time to complete each part was recorded. Each part’s score represents the time required to complete the task.

#### Neuroimaging Procedures

Participants completed the BeST Memory Task while inside the 3T Siemens MRI Scanner at the Magnetic Resonance Imaging and Spectroscopy Center (MRISC) of the University of Kentucky using a 32-channel head coil. High-resolution anatomic images (20 minutes 3D MPRAGE) were acquired using a rapid gradient echo acquisition sequence (acquisition matrix 256 x 256 x 176, isotropic 1 mm voxels, field of view 256 mm, repetition time 2530 ms, echo time 2.26 ms). With fMRI and as part of a larger study, we analyzed the volume of multiple regions of interest, including the hippocampus, amygdala, dorsolateral prefrontal cortex (DLPFC), frontal eye field (FEF), precuneus, inferior parietal lobe (IPL), insula, and the fusiform. The volumes of each region of interest were normalized to the total intracranial volume. CSF samples of Aß_42_ and p-Tau_181_ were taken from all participants as described in the Jiang et al. (2016) study. In this study, we present findings on precuneus volume.

### Data analysis

Descriptive statistics was performed to calculate participants’ demographic and clinical characteristics. Pearson *r* correlation analyses were performed on all 45 participants and in separate sex categories (men and women). Additionally, Pearson *r* correlation analyses were performed on the neuropsychological and fMRI data with age and education controlled to account for potential confounding variables.

## Results

### Participant demographics

Participants’ demographic and clinical characteristics are summarized in Table 1. The mean participant age was 77.05 (±16.54), with a range of 59-93 years. The mean number of years of formal education was 16.54 (±2.46), and 55.6% of the subjects were women. Age (*p*= 0.870) and level of education (*p*=0.412) did not differ significantly between men and women.

**Table 1:** Demographic and clinical characteristics (N = 45)

| **Variable** | **All participants** | **Men**<br>***n*=20** | **Women**<br>***n*=25** | ***p* value** |
| --- | --- | --- | --- | --- |
|  | ***M (SD)/Yes*** | ***M (SD)/Yes*** | ***M (SD)/Yes*** |  |
| Age | 77.05 (7.48) | 76.83 (8.08) | 77.24 (7.11) | 0.870 |
| Education | 16.54 (2.46) | 16.89 (2.30) | 16.24 (2.61) | 0.413 |
| Trail A | 35.68 (15.20) | 33.28 (12.08) | 37.85 (17.57) | 0.353 |
| Trail B | 90.76 (54.19) | 85.67 (60.11) | 95.35 (49.39) | 0.593 |
| APOE | 7 | 5 | 2 |  |
| A-Beta42 | 265.88 (81.26) | 264.26 (81.04) | 267.17 (83.15) | 0.909 |
| p-Tau181P | 23.47 (7.68) | 23.42 (6.86) | 23.50 (8.42) | 0.973 |

### Sex differences in memory performances

There were statistically significant differences between men and women in some aspects of memory performance (Figure 2). Men and women differed in their reaction time to distractor images (*p*=0.023), their reaction time to correct responses (p=0.002), and their false alarm reaction rate (*p*=0.044). Women responded slower to distractor images than men and produced correct responses. Also, women were slower in reacting to false alarms compared to men. Thus, showing higher accuracy in memory tasks than men. See Table 2 for additional information.

**Figure 2a.**
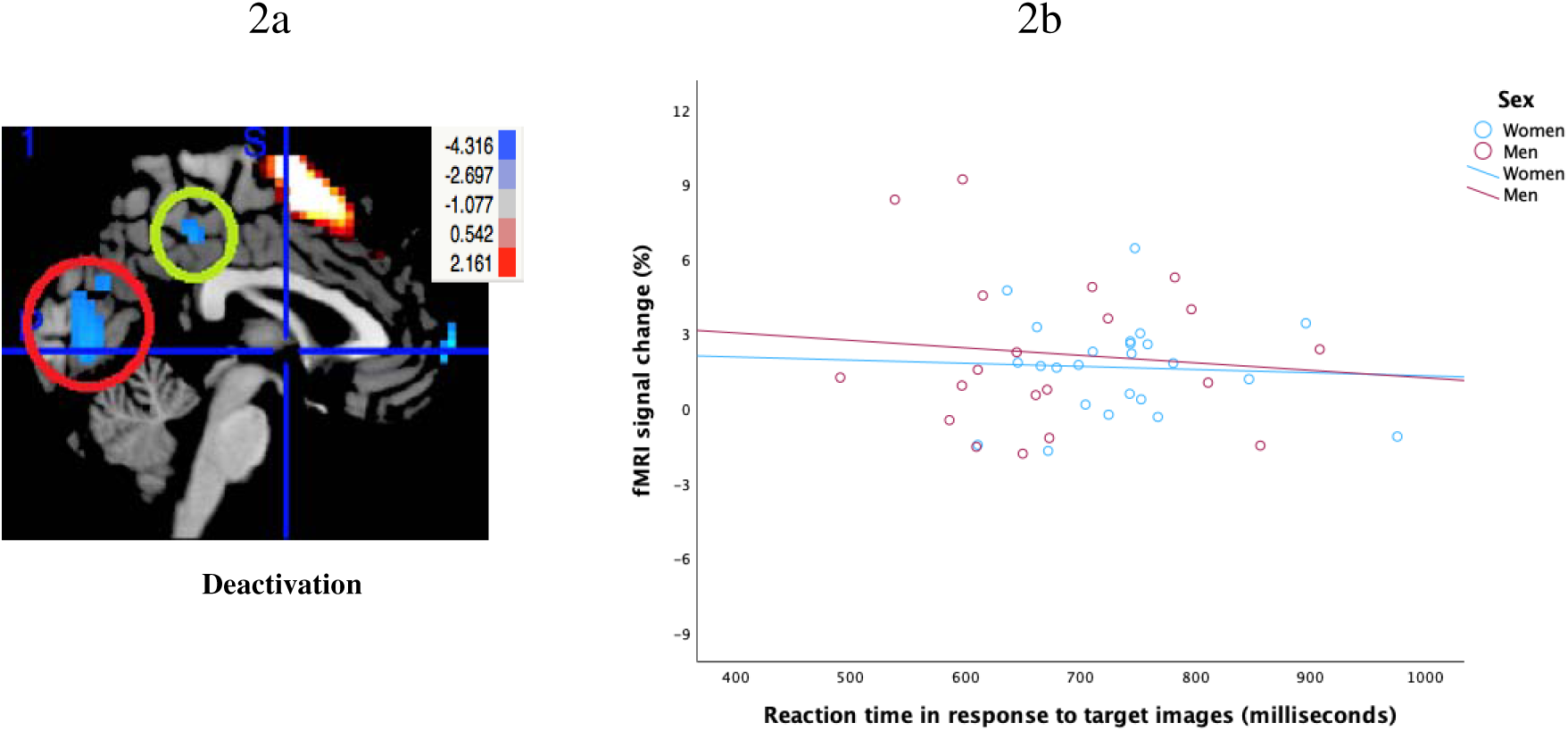
shows the deactivation of fMRI-BOLD signals in the precuneus. Figure 2b shows the percentage of fMRI signal change across reaction time in response to target images for men and women.

**Table 2:** Descriptives result of memory performances.

| **Variables** | **All participants** | **Men** | **Women** | **p value** |
| --- | --- | --- | --- | --- |
|  | *M (SD)* | *n=20*<br>*M (SD)* | *n=25*<br>*M (SD)* |  |
| RT1 | 708.76 (98.09) | 676.51 (107.96) | 735.64 (81.86) | 0.052* |
| RT2 | 745.27 (93.21) | 709.40 (104.66) | 775.15 (71.73) | 0.023* |
| CR_RT | 724.70 (85.69) | 679.46 (83.70) | 763.47 (67.64) | 0.002** |
| FA_RT | 728.85 (193.67) | 656.83 (165.76) | 792.39 (198.68) | 0.044* |
| FA_Rate | 0.04 (0.037) | 0.04 (0.03) | 0.04 (0.04) | 0.810 |
| CR_Rate | 0.91 (0.11) | 0.92 (0.12) | 0.91 (0.11) | 0.788 |
| AC1 | 0.90 (0.14) | 0.88 (0.21) | 0.91 (0.06) | 0.530 |
| AC2 | 0.95 (0.08) | 0.94 (0.11) | 0.96 (0.04) | 0.424 |
| CR_N | 88.49 (13.23) | 90.11 (11.37) | 87.10 (14.77) | 0.476 |
| FA_N | 4.63 (3.56) | 4.40 (3.25) | 4.82 (3.91) | 0.740 |

*Note:* \*= $p < .05$ ; \*\* $p < .01$ . RT1= Reaction time in response to target images, RT2=Reaction time in response to distractor images, AC1= Accuracy in response to target images, AC2= Accuracy in response to distractor images, CR\_N=correct response number, CR\_R=correct response rate, CR\_RT= Reaction times of correct responses, FA\_N=false alarm number, FA\_Rate=FA\_RT=false alarm reaction time.

### Precuneus deactivation correlates with faster reaction time

We examined the relationship between memory tasks and resting-state functional brain connectivity. Figure 2a shows the deactivation of the precuneus in milliseconds, while Figure 2b shows the relationship between fMRI signal change during precuneus deactivation and reaction time to in response to target images in milliseconds. We found that there was a statistically significant correlation between precuneus deactivation and reaction time in the sample (*p*=0.013).

### Precuneus volumes display negative correlation with reaction time in women but not in men

Figure 4a shows the deactivation patterns of the posterior precuneus in the participants during the working memory task. The results (see Table 3) show that in women, volumes in the left precuneus displayed a significant negative correlation with reaction time in response to target images (*r* = -0.426; *p* < 0.05) and distractor images (*r* = -0.549; *p* < 0.01). As volumes in the left precuneus decreased, reaction time in response to both target and distractor images increased. There was no significant relationship between left precuneus volumes and reaction time in response to target or distractor images in men. The results also show volumes in the left precuneus displayed a significant negative correlation with reaction time to correct responses in women (*r* = -0.580; *p* < 0.01). As the volumes in the left precuneus increased, reaction time to correct responses decreased.

**Figure 3.**
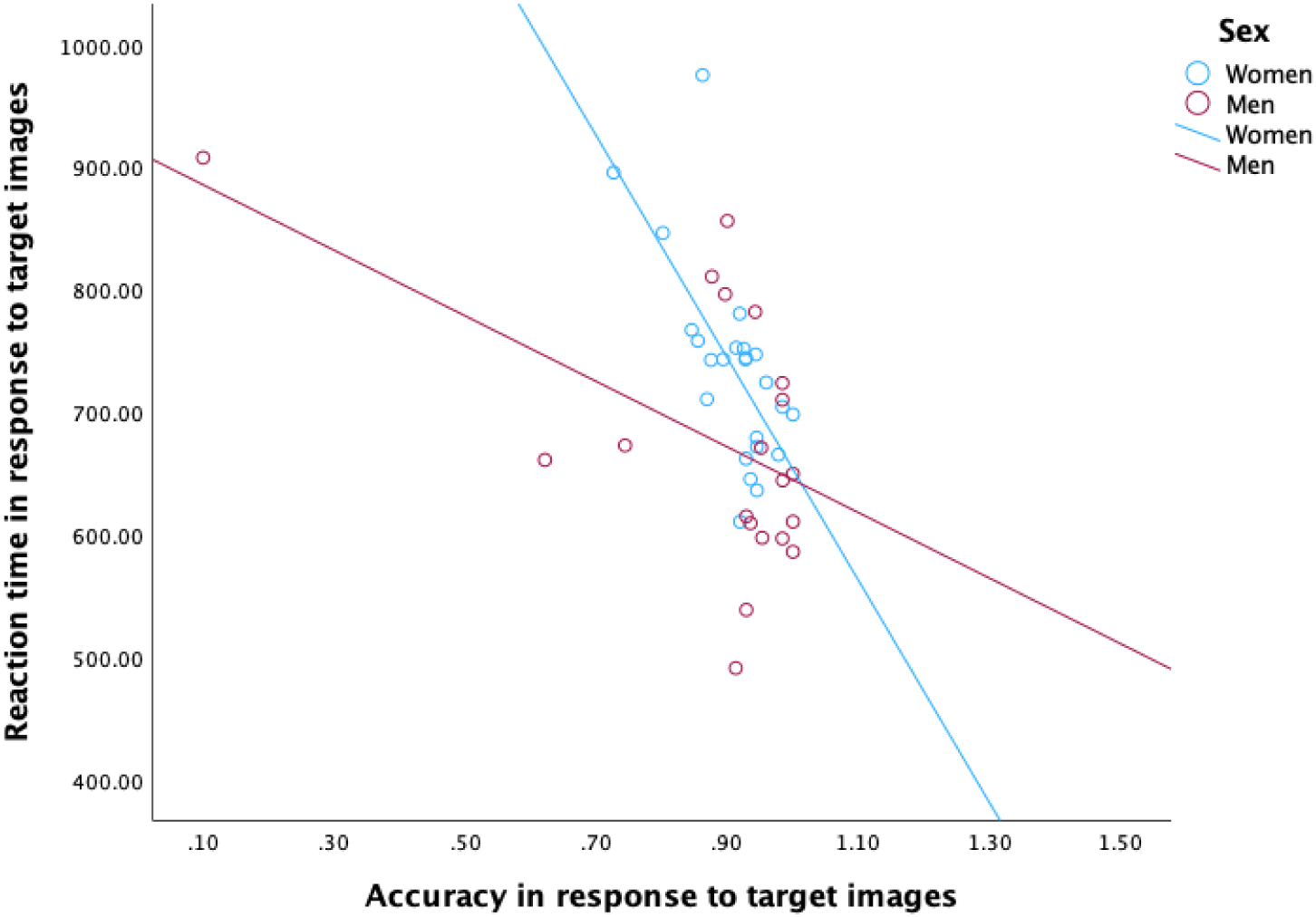
Sex difference in accuracy and interaction trade-off of reaction time in response to target images for men and women.

**Figure 4a.**
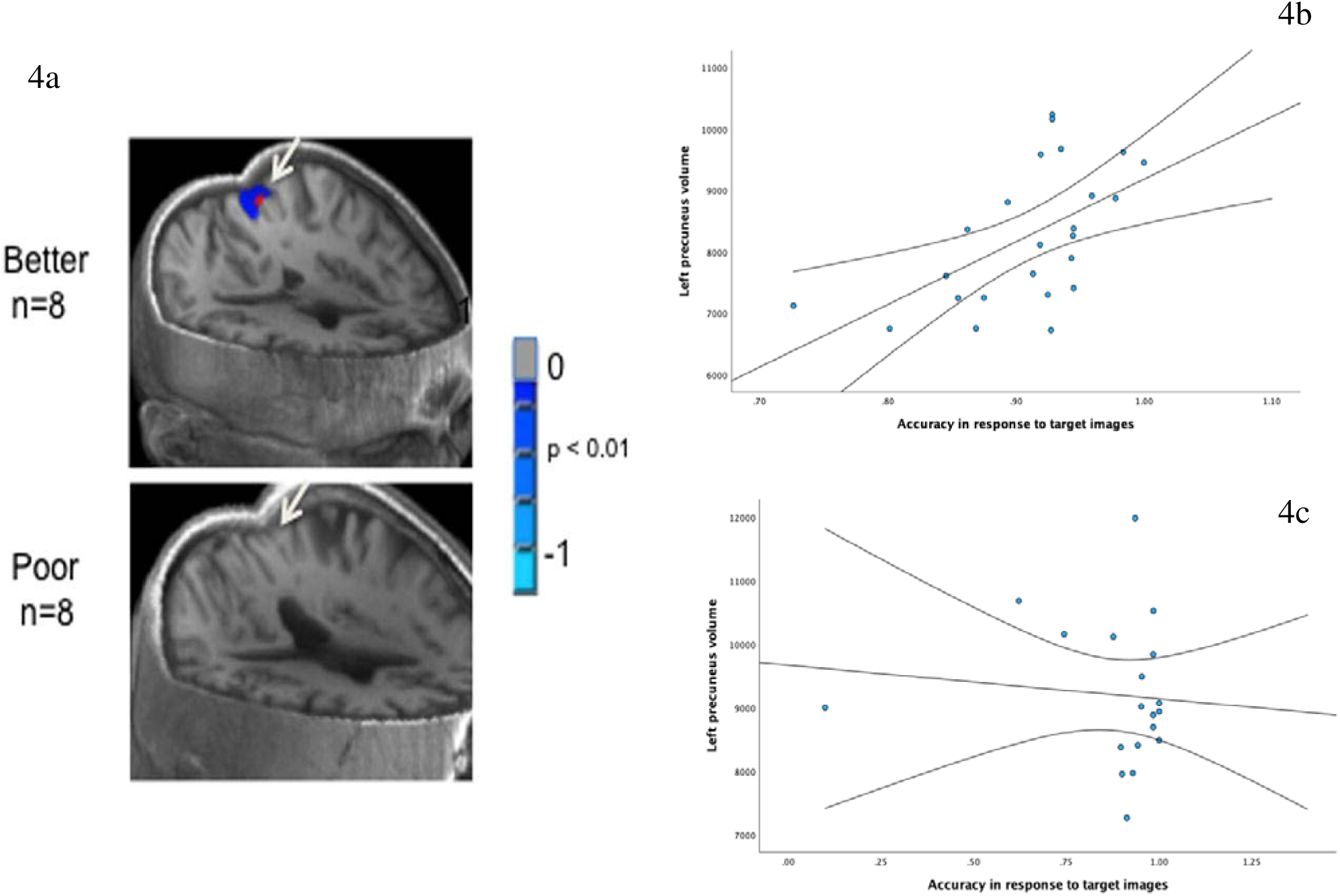
shows the top eight versus bottom eight performers in a delayed match-to-sample working memory task. Better performers on the working memory task showed deactivation of posterior precuneus/cingulate (top) during the working memory task, whereas poor performers failed to show the deactivation pattern (bottom). Figure 4b shows the correlation between Left Precuneus volume and accuracy in response to target images in women. Figure 4c shows the correlation between Left Precuneus volume and accuracy in response to target images in men.

**Table 3:**
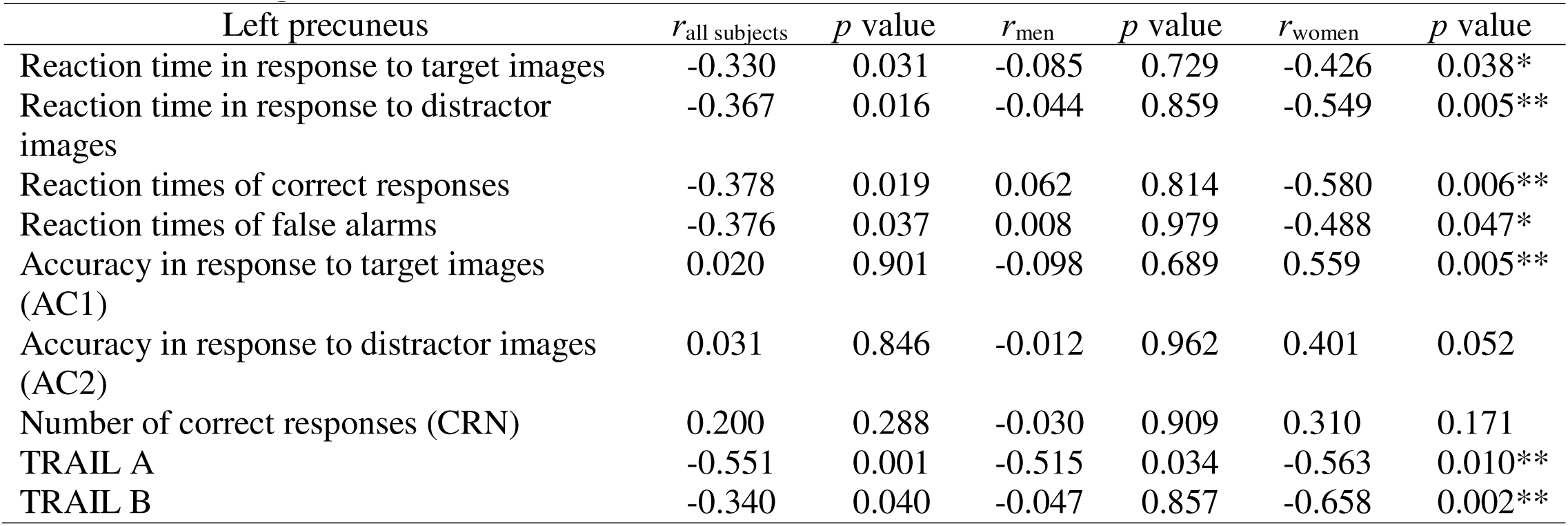
Correlations between left precuneus volumes, and reaction time, accuracy of BeST task and trailmaking task in men and women.

| Left precuneus | $r_{\text{all subjects}}$ | $p$ value | $r_{\text{men}}$ | $p$ value | $r_{\text{women}}$ | $p$ value |
| --- | --- | --- | --- | --- | --- | --- |
| Reaction time in response to target images | -0.330 | 0.031 | -0.085 | 0.729 | -0.426 | 0.038* |
| Reaction time in response to distractor images | -0.367 | 0.016 | -0.044 | 0.859 | -0.549 | 0.005** |
| Reaction times of correct responses | -0.378 | 0.019 | 0.062 | 0.814 | -0.580 | 0.006** |
| Reaction times of false alarms | -0.376 | 0.037 | 0.008 | 0.979 | -0.488 | 0.047* |
| Accuracy in response to target images (AC1) | 0.020 | 0.901 | -0.098 | 0.689 | 0.559 | 0.005** |
| Accuracy in response to distractor images (AC2) | 0.031 | 0.846 | -0.012 | 0.962 | 0.401 | 0.052 |
| Number of correct responses (CRN) | 0.200 | 0.288 | -0.030 | 0.909 | 0.310 | 0.171 |
| TRAIL A | -0.551 | 0.001 | -0.515 | 0.034 | -0.563 | 0.010** |
| TRAIL B | -0.340 | 0.040 | -0.047 | 0.857 | -0.658 | 0.002** |

There was no significant relationship between left precuneus volumes and reaction time to correct responses in men. Additionally, the results show that volumes in the left precuneus displayed a significant negative correlation with reaction time to false alarms (*r* = -0.488; *p* < 0.05) in women. As volumes in the left precuneus increased, reaction time to false alarms decreased. There was no significant relationship between left precuneus volumes and reaction time to correct responses in men. Further, we found that volumes in the left precuneus displayed significant positive correlation with accuracy in response to target images (r = 0.559; p < 0.01; Figure 4b) and distractor images (r = 0.401; p = 0.052) in women. As the volumes in the left precuneus increased, accuracy in response to both target and distractor images also increased. There was no significant relationship between left precuneus volumes and accuracy in response to target (Figure 4c) or distractor images in men. Also, volumes in the left precuneus displayed no significant correlation with the total number of correct responses in both men and women. See Table 3 for detailed information.

Table 3 also shows that volumes in the left precuneus displayed significant negative correlation with both Trail Making task A (*r* = -0.563 p = 0.010) and B (*r* = -0.658, p < 0.01) in women. As the volumes in the left precuneus increased, the time needed to complete trailmaking tasks decreased. Also, volumes in the left precuneus displayed a significant negative correlation with Trail Making task A in men (*r* = -0.515 p < 0.05). There was no significant relationship between left precuneus volumes and time to complete Trail Making task B in men.

## Discussion

As the number of people at risk of developing AD increases, the characterization of sex differences in AD risk is important for the direction of future research, precision diagnoses, and therapeutic interventions (Mielke 2020; Nebel et al. 2018). We aimed to examine potential sex differences in the precuneus volume and compared the participants’ memory performance on memory and neuropsychological tasks. By studying sex differences in healthy individuals, researchers may gain insights into how these differences contribute to AD risk, progression, and treatment response while addressing the complex interplay between biological factors and memory-related outcomes.

We hypothesized that sex differences would have differential effects on brain volumes and network-associated working memory performance in cognitively normal older adults. We found that in women, measures of working memory performance correlated consistently with volume in the left precuneus. While men were faster in reaction time, women were better in accuracy. These relationships were consistent and in expected directions given units of measure (e.g. negative correlation between left precuneus volume and trailmaking task score, but positive correlation between left precuneus volume and accuracy). Our study supports prior findings that pre-AD pathology differs in women versus men (Medeiros & Silva et al., 2019; Samson et al., 2023; Williamson et al., 2022). By identifying that women exhibit slower reaction time, but better accuracy compared to men in a memory task, this study contributes to a deeper understanding of the neural mechanisms underlying cognitive processes and highlights the need for further exploration of sex-related differences in cognition.

Atrophy of brain structures has been continuously recognized as an early and key feature of AD (Jiang et al., 2016; Richter et al., 2022; Rusinek et al., 2004; Williamson et al., 2022). Since the precuneus plays a critical role in episodic memory retrieval, early changes in the precuneus can signal memory impairment—a precursor for AD. Studies show that AD pathology tends to follow a characteristic pattern of progression, starting in the MTL and spreading to other brain regions over time (Rusinek et al. 2004). This temporal progression aligns with the clinical observation that memory deficits are often among the earliest symptoms of the disease (Aël Chetelat & Baron, 2003). Similarly, autopsy studies of individuals with AD consistently reveal significant atrophy in brain structures (de Flores et al., 2020; Nedelska et al., 2015; Whitwell et al., 2008). In line with the current study, advanced neuroimaging techniques, such as fMRI, allow us to visualize structural changes in the brains and the associated cognitive and neurological conditions.

Our findings point to the potential of reaction time serving as a surrogate marker for fMRI in predicting cognitive decline, particularly when considering sex differences. Research has shown that slower reaction times are associated with cognitive impairment and decline, as they may reflect deficits in processing speed, attention, and executive function, which are early indicators of cognitive decline (Haynes et al. 2017; Velichkovsky et al. 2020). We showed that women tend to have slower reaction times compared to men, but they may compensate with better accuracy in memory tasks. Therefore, monitoring changes in reaction time, especially in conjunction with fMRI measures of brain function, could provide valuable insights into sex-specific patterns of cognitive decline. By integrating reaction time as a surrogate marker with fMRI data, researchers can better understand the underlying neural mechanisms of cognitive decline and potentially identify sex-specific biomarkers for early detection and targeted interventions.

In addition to adding to the evidence that there are sex differences in AD risk factors, our findings have the potential to significantly inform targeted interventions aimed at preserving cognitive function in affected individuals. One strength of the present study is that many working memory measures were collected, including reaction time, accuracy, and false alarm. Other memory tasks were also utilized, including animal naming, vegetable naming, and trail making tasks. This variety of data allowed for the analysis of multiple aspects of working memory. Although many studies have reported developmental differences in parietal lobe between boys and girls, very little is known in brain and cognitive aging related to the brain structure. The current study is unique in terms of sex-specific working memory performances in bilateral precuneus function in aging population.

Some limitations are worth noting. The present study has a relatively small sample size. A larger sample size may yield findings that are different from those in the current study. The sample is all non-Hispanic White; therefore, our findings cannot be generalized to racially diverse older Americans. Additionally, though our findings focus on the default mode network (DMN), initial identification of regions of interest and measurements of those regions focused on the medial temporal and frontal lobes as those are often sites of first pathology in AD. Though those regions showed variable relationships with working memory measures, no relationship was as consistent and predictable as that with the precuneus. In the future, we hope to focus these measurements on the DMN to further characterize its relationship with working memory in preclinical older adults and how this relationship may differ between men and women. Furthermore, we hope to characterize how this relationship changes over time. The present study examined only cognitively normal older adults, but future studies should investigate these differences in adults with clinical AD.

## Conclusion

The current study shows that by studying sex differences in brain volumes, researchers may gain insights into how sex differences contribute to memory impairment and AD risk. Identifying sex-specific factors that influence memory can guide the development of targeted preventive measures and interventions. This is particularly relevant as researchers explore ways to promote cognitive health and resilience against age-related memory decline. Since women tend to have a higher prevalence of certain neurodegenerative disorders, understanding sex differences in memory-related brain regions can inform policies, healthcare strategies, and resources for managing and preventing memory-related disorders.

## Acknowledgement

We thank A. Anderson and E Wallis for volume analysis, the human participants from the University of Kentucky Alzheimer’s Disease Research (UK-ADRC) Cohort, and members of the Aging Brain and Cognition (ABC) laboratory.

## Authors’ contributions

DP, GJ, and YJ contributed to the study’s conception and design. Literature search was performed by DE, SS, AB, and LB. DP, AB, LB, and YJ conducted data analyses of behavioral and MRI data, with additional statistical analysis from DE. DE, SS, and YJ contributed to the writing of the manuscript.

